# Effects of enalapril and paricalcitol treatment on diabetic nephropathy and renal expressions of TNF-α, P53, Caspase-3 and Bcl-2 in STZ-induced diabetic rats

**DOI:** 10.1101/577106

**Authors:** Osama M. Ahmed, Tarek M. Ali, Mohamed A. Abdel Gaid, Ahmed A. Elberry

## Abstract

This study aimed to assess the renopreventive effect of the angiotensin converting enzyme inhibitor (ACEI), enalapril, and/or vitamin D receptor (VDR) activator, paricalcitol, on streptozotocin (STZ) diabetes-induced nephropathy and to elucidate the mechanisms of action through investigation of the effects on renal oxidative stress, antioxidant defense system and expressions of TNF-α, P53, caspase-3, and Bcl-2. Diabetes mellitus was induced in fasting male Wistar rats by single intraperitoneal injection of STZ (45 mg /kg b.w.) dissolved in citrate buffer pH 4.5. Ten days after STZ injection, the diabetic rats were treated with enalapril (25 mg/l of drinking water) and/or paricalcitol (8 µg/kg b.w. *per os*) dissolved in 5% DMSO daily for 4 weeks. The obtained data revealed that the treatment of diabetic Wistar rats with enalapril and/or paricalcitol led to a significant decrease in the elevated serum urea, uric acid, creatinine and sodium, potassium levels; thereby reflecting improvement of the impaired kidney function. The deteriorated kidney lipid peroxidation, GSH content and GST and catalase activities in diabetic rats were significantly ameliorated as a result of treatment with enalapril and/or paricalcitol. The elevated fasting and post-prandial serum glucose levels and the lowered serum insulin and C-peptide levels were also improved. Moreover, the treatment of diabetic rats successfully prevented the diabetes-induced histopathological deleterious changes of kidney and islets of Langerhans of pancreas. In association, the immunohistochemically detected pro-inflammatory cytokine TNF-α and apoptotic mediators P53 and caspase-3 were remarkably decreased in kidney of diabetic rats as a result of treatment, while the expression of anti-apoptotic protein Bcl-2 was increased. Based on these findings, it can be concluded that enalapril and paricalcitol can prevent STZ diabetes-induced nephropathy though amelioration of the glycemic state and antioxidant defense system together with the suppression of oxidative stress, inflammation and apoptosis.

## Introduction

Diabetic nephropathy (DN), a complicated microvascular disease associated with diabetes mellitus (DM), is a significant cause of chronic kidney disease (CKD) [1]. In a type 1 diabetes mellitus (T1DM) population with a mean age of 44 years and duration of diabetes of 17 years, up to 21.0% of patients developed CKD [2]. Similarly, DN occurs in 20% to 40% of all cases of type 2 diabetes mellitus (T2DM) [3]. Approximately 14.5% mortality was recorded among people with DM (aged 20 to 79 years) [4]. This increased risk may be essentially driven by kidney disease which may be correlated with the presence of atherosclerosis or diabetes-associated glomerular damage [5] probably with the incidence of severe and interstitial inflammation [6].

The prevention of activated renin-angiotensin-aldosterone system (RAAS) in diabetic kidney has a vital role in treatment of DN [7]. Drugs that block RASS pathway are effective in reducing nephropathy progression in diabetic patients and delaying cardiovascular and renal morbidity and mortality [8].

Enalapril is one of angiotensin converting enzyme inhibitors (ACEIs), a class of antihypertensive medications, that have been shown to be of greater benefit in preventing RAAS, reducing DN, abating progression of renal failure, restoring glomerular hyperperfusion and hyperfiltration, improving glomerular barrier function, and reducing the non-hemodynamic effects of angiotensin II and aldosterone [9,10]. Moreover, Vermes *et al*. [11] reported that enalapril markedly reduces the risk of developing diabetes in patients with left ventricular systolic dysfunction by elevation of insulin sensitivity, skeletal muscle glucose uptake, and pancreatic blood flow.

Paricalcitol (19-nor-1, 25-dihydroxyvitamin D2), an active non-hypercalcemic selective vitamin D analogue, was reported to inhibit activated RAAS by decreasing renin, renin receptor, angiotensinogen and angiotensin type 1 receptor [12]. Studies in experimental nephropathy animal models have demonstrated that paricalcitol improves glomerular degeneration and tubular damage [13]. Clinically, Aperis *et al*. [14] demonstrated an average 32.9 % reduction of proteinuria in 19 patients treated with 1-2 μg daily paricalcitol with 74 % of patients respond, especially in diabetic nephropathy. Izquierdo *et al*. [15] found that paricalcitol has antioxidant effects as it decreases serum malondialdhyde (MDA) levels and increases serum reduced glutathione (GSH) and thioredoxin (TRX) levels, as well as superoxide dismutase (SOD) and catalase (CAT) activities in hemodialysis patients.

Therefore, this study was conducted to assess the renopreventive effect of the angiotensin converting enzyme inhibitor, ACEI, enalapril, and vitamin D receptor (VDR) activator, paricalcitol, on streptozotocin (STZ)-induced nephropathy through investigating effect on renal oxidative stress, antioxidant defense system and expressions of tumor necrosis factor-α (TNF-α), caspase-3, B cell lymphoma-2 (Bcl-2), and protein 53 (P53) genes.

## Materials and Methods

### Experimental Animals

Male albino rats of Wistar strain weighing about 100-130 grams were used in this study. After two weeks of adaptation period, the animals were housed in clean polypropylene cages and maintained in an air-conditioned animal house at temperature (20-25oC) with natural alternating light and dark cycles. The animals were supplemented with standard pellet diets and water *ad libitum*. All experimental animal procedures are in accordance with the guidelines of the Experimental Animal Ethics Committee of Faculty of Science, Beni-Suef University, Beni-Suef, Egypt (Ethical Approval Number: BSU/FS/2016/11).

### Induction of Diabetes Mellitus

Diabetes mellitus was induced in male Wistar rats by a single intraperitoneal injection of streptozotocin (STZ) (Sigma, St. Louis, MO, USA) at dose level of 45 mg /kg b.w. dissolved in citrate buffer (pH 4.5) [16]. Ten days after STZ injection, rats were deprived of food and water overnight (10–12 h) and blood samples were taken from lateral tail vein at fasting state and 2 hours (hr) of oral glucose loading (3 g/kg b.w.). Serum glucose concentration was measured. Rats with a 2-hr serum glucose level ranging from 180 to 300 mg/dl were considered as mild diabetic while those outside this range were excluded.

### Animal grouping

After induction of diabetes mellitus, the rats were allocated into the five groups (6 rats for each group): Group 1 (Normal group) was intraperitoneally administered the equivalent volume of the vehicle (5% dimethyl sulphoxide [DMSO]). The rats of Group 2 (Diabetic control group) were diabetic rats that were intraperitoneally administered the equivalent volume of the vehicle (5% dimethyl sulphoxide [DMSO]) for 4 weeks. Group 3 (Diabetic group treated with enalapril) included diabetic rats that were treated with enalapril in drinking water at concentration 25 mg enalapril/l of drinking water for 4 weeks; this group was intraperitoneally administered the equivalent volume of 5% DMSO. Group 4 (Diabetic group treated with paricalcitol) consisted of diabetic rats that were intraperitoneally injected with paricalcitol at dose level of 8 µg/kg b. w. dissolved in 5% DMSO daily for 4 weeks. Group 5 ((Diabetic group treated with enalapril and paricalcitol) was composed of diabetic rats that were treated with enalapril in drinking water (25 mg enalapril/l) and were also intraperitoneally injected with pariclacitol at dose level of 8 µg/kg b. w. dissolved in 5% DMSO daily for 4 weeks.

### Blood and tissue sampling

At the end of the experimental period, animals were sacrificed under anesthesia. Blood samples from jugular vein were collected. The blood was left to coagulate at room temperature and then centrifuged at 3000 r.p.m. for 30 minutes. The clear nonhaemolysed supernatant sera were quickly removed and kept at −30 °C pending their use in analysis of various biochemical parameters related to kidney functions.

Kidneys of each animal were excised and then one kidney of each rat was fixed in neutral buffer formalin for histopathological and immunohistochemical studies. A 0.5 g from other kidney of each animal was homogenized in 5 ml 0.9% NaCl (10%w/v) using Teflon homogenizer (Glas-Col, Terre Haute, USA). The obtained homogenates was centrifuged at 3000 r.p.m. for 10 minutes and the homogenate supernatants were aspirated and kept in deep freezer at −30°C to be used later for measurements of markers of oxidative stress and antioxidant defense system.

### Biochemical investigations

Serum urea level was determined using kits obtained from BIOMED Diagnostic (EGY-CHEM for Lab Technology) Bader city, Cairo, Egypt. Serum uric acid was determined using reagent kits purchased from Spinreact, S.A.U. (SPAIN). Creatinine was detected using reagent kits obtained from Diamond Diagnostic Chemical Company (Egypt). Measurement of sodium and potassium ions were carried out using reagent kits purchased from Spectrum Company for Biotechnology, Obour City, Cairo, Egypt. Serum glucose levels at fasting state and after 2 hours of oral glucose loading were determined by using commercial diagnostic kit obtained from Randox Laboratories, UK. Serum insulin and C-peptide levels were assayed using enzyme-linked immunosorbent assay kits purchased from Linco Research, St. Charles, MO, USA according to manufacturer’s instruction. Kidney GSH level, lipid peroxidation (LPO) represented by MDA level, glutathione-S-transferase (GST) activity, and catalase (CAT) activity were determined according to the chemical methods of Beutler *et al*. [17], Preuss *et al*. [18], Mannervik and Gutenberg [19] and Cohen *et al*. [20] respectively.

### Histological and immunohistochemal investigations

Fixed kidneys were transferred to National Cancer Institute, Cairo University, Egypt for blocking in wax, sectioning and staining with hematoxylin and eosin according to the method of Banchroft *et al*. [21]. Then, the prepared stained sections were examined to detect the histological changes.

Immunohistochemical techniques for TNF-α, P53, caspase-3 and Bcl-2 by using 3 μm thickness liver sections mounted on positive glass slides according to the methods of Hussein and Ahmed [22] respectively.

### Statistical analysis

The data were analysed using the one-way analysis of variance (ANOVA) (PC-STAT, 1985, University of Georgia, USA) [23] followed by LSD analysis to compare various groups with each other. Results were expressed as mean ± standard error (SE). F-probability obtained from one-way ANOVA, expresses the effect between groups.

## Results

Serum urea, uric acid and creatinine levels were significantly (p<0.01; LSD) elevated in diabetic control group while this elevation was remarkably decreased in diabetic treated groups with F-probabilities P<0.001, P<0.05 and P<0.01 respectively (Table 1). The treatment with enalapril produced the most potent effects in decreasing the elevated serum urea, uric acids and creatinine levels recording percentage changes of −49.36, −39.72 and −41.50% respectively. Both serum sodium and potassium levels were significantly increased (p<0.01; LSD) in diabetic control rats as compared with normal control; the recorded percentage changes were 3.59 and 22.04% respectively. The treatment of diabetic rats with enalapril, paricalcitol and their combinations significantly normalized the elevated serum sodium and potassium levels when compared with diabetic control with F-probabilities P<0.001 and P<0.05 respectively (Table 2). The elevated kidney LPO represented by MDA level was normalized by the treatment with enalapril, paricalcitol and their combination recording percentage changes of −40.11, −44.60 and −36.57 respectively; thereby, paricalcitol appeared to be the most potent (Table 3). The lowered kidney GSH content as well as GST and catalase activities in diabetic rats were significantly increased as a result of treatment with enalapril, paricalcitol and their combination with F-probabilities of P<0.001, P<0.01 and P<0.001 respectively (tables 3 and 4). While the effect of co-treatment with enalapril and paricalcitol was the most potent on GSH content (37.49%), paricalcitol appeared to be the most effective on GST activity (43.64%) and enalapril seemed to be the most effective on catalase activity (40.70%) (Tables 3 and 4).

**Table 1:** Effects of enalapril and paricalcitol on serum urea, uric acid and creatinine levels in diabetic rats.

| Group \ Parameter | Urea (mg/dl) | % change | Uric acid (mg/dl) | % change | Creatinine (mg/dl) | % change |
| --- | --- | --- | --- | --- | --- | --- |
| Normal | 29.17 ± 2.29<sup>d</sup> | - | 1.31 ± 0.132<sup>b</sup> | - | 0.65 ± 0.004<sup>b</sup> | - |
| Diabetic control | 83.52 ± 9.79<sup>a</sup> | 186.32 | 2.19 ± 0.295<sup>a</sup> | 67.17 | 1.06 ± 0.138<sup>a</sup> | 63.07 |
| Diabetic treated with Enalapril | 42.29 ± 1.54<sup>cd</sup> | -49.36 | 1.32 ± 0.181<sup>b</sup> | -39.72 | 0.62 ± 0.003<sup>b</sup> | -41.50 |
| Diabetic treated with Paricalcitol | 65.56 ± 5.15<sup>b</sup> | -21.50 | 1.52 ± 0.140<sup>b</sup> | -30.59 | 0.78 ± 0.082<sup>b</sup> | -26.41 |
| Diabetic treated with Enalapril and Paricalcitol | 55.67 ± 5.17<sup>bc</sup> | -33.34 | 1.74 ± 0.230<sup>ab</sup> | -20.54 | 0.70 ± 0.074<sup>b</sup> | -33.69 |
| F-probability | $P<0.001$ | | $P<0.05$ | | $P<0.01$ | |
| LSD at 5% level | 16.322 |  | 0.598 |  | 0.240 |  |
| LSD at 1% level | 22.082 |  | 0.809 |  | 0.325 |  |

- Data are expressed as mean ± standard error. Number of detected samples in each group is six. - Means, which share the same superscript symbol(s) are not significantly different. - Percentage changes were calculated by comparing diabetic control group with normal control group and diabetic treated groups with diabetic control group.

**Table 2:** Effects of enalapril and paricalcitol on serum sodium and potassium levels in diabetic rats.

| Parameter<br>Group | Sodium<br>(mg/dl) | %<br>change | Potassium<br>(mg/dl) | %<br>change |
| --- | --- | --- | --- | --- |
| Normal | 149.19 ± 0.38<sup>c</sup> | - | 4.90 ± 0.21<sup>b</sup> | - |
| Diabetic control | 154.55 ± 0.84<sup>a</sup> | 3.59 | 5.98 ± 0.26<sup>a</sup> | 22.04 |
| Diabetic treated with Enalapril | 150.27 ± 0.12<sup>bc</sup> | -2.76 | 5.16 ± 0.04<sup>b</sup> | -13.71 |
| Diabetic treated with Paricalcitol | 151.84 ± 0.35<sup>b</sup> | -1.75 | 5.23 ± 0.35<sup>b</sup> | -12.54 |
| Diabetic treated with Enalapril and Paricalcitol | 151.59 ± 0.74<sup>b</sup> | -1.91 | 4.83 ± 0.15<sup>b</sup> | -19.23 |
| F-probability | P<0.001 |  | P<0.05 |  |
| LSD at 5% level | 1.627 |  | 0.668 |  |
| LSD at 1% level | 2.202 |  | 0.904 |  |

- Data are expressed as mean ± standard error. Number of detected samples in each group is six.
- Means, which share the same superscript symbol(s) are not significantly different.
- Percentage changes were calculated by comparing diabetic control group with normal control group and diabetic treated groups with diabetic control group.

**Table 3:** Effects of enalapril and paricalcitol on kidney GSH content and LPO in diabetic rats.

| Parameter<br>Group | LPO<br>(nmole MDA/100<br>mg tissue/hr) | %<br>change | GSH<br>(nmole/100<br>mg tissue) | %<br>change |
| --- | --- | --- | --- | --- |
| Normal | 20.00 ± 1.17<sup>b</sup> | - | 73.90 ± 5.08<sup>a</sup> | - |
| Diabetic control | 32.26 ± 2.75<sup>a</sup> | 61.30 | 52.06 ± 0.73<sup>b</sup> | -29.55 |
| Diabetic treated with Enalapril | 19.32 ± 3.19<sup>b</sup> | -40.11 | 70.15 ± 7.40<sup>a</sup> | 34.94 |
| Diabetic treated with Paricalcitol | 17.87 ± 1.04<sup>b</sup> | -44.60 | 48.94 ± 0.63<sup>b</sup> | -5.99 |
| Diabetic treated with Enalapril and Paricalcitol | 20.46 ± 1.60<sup>b</sup> | -36.57 | 71.58 ± 3.47<sup>a</sup> | 37.49 |
| F-probability | P< 0.001 |  | P< 0.001 |  |
| LSD at 5% level | 6.230 |  | 12.609 |  |
| LSD at 1% level | 8.429 |  | 17.059 |  |

- Data are expressed as mean ± standard error. Number of detected samples in each group is six.
- Means, which share the same superscript symbol(s) are not significantly different.
- Percentage changes were calculated by comparing diabetic control group with normal control group and diabetic treated groups with diabetic control group.

**Table 4:**
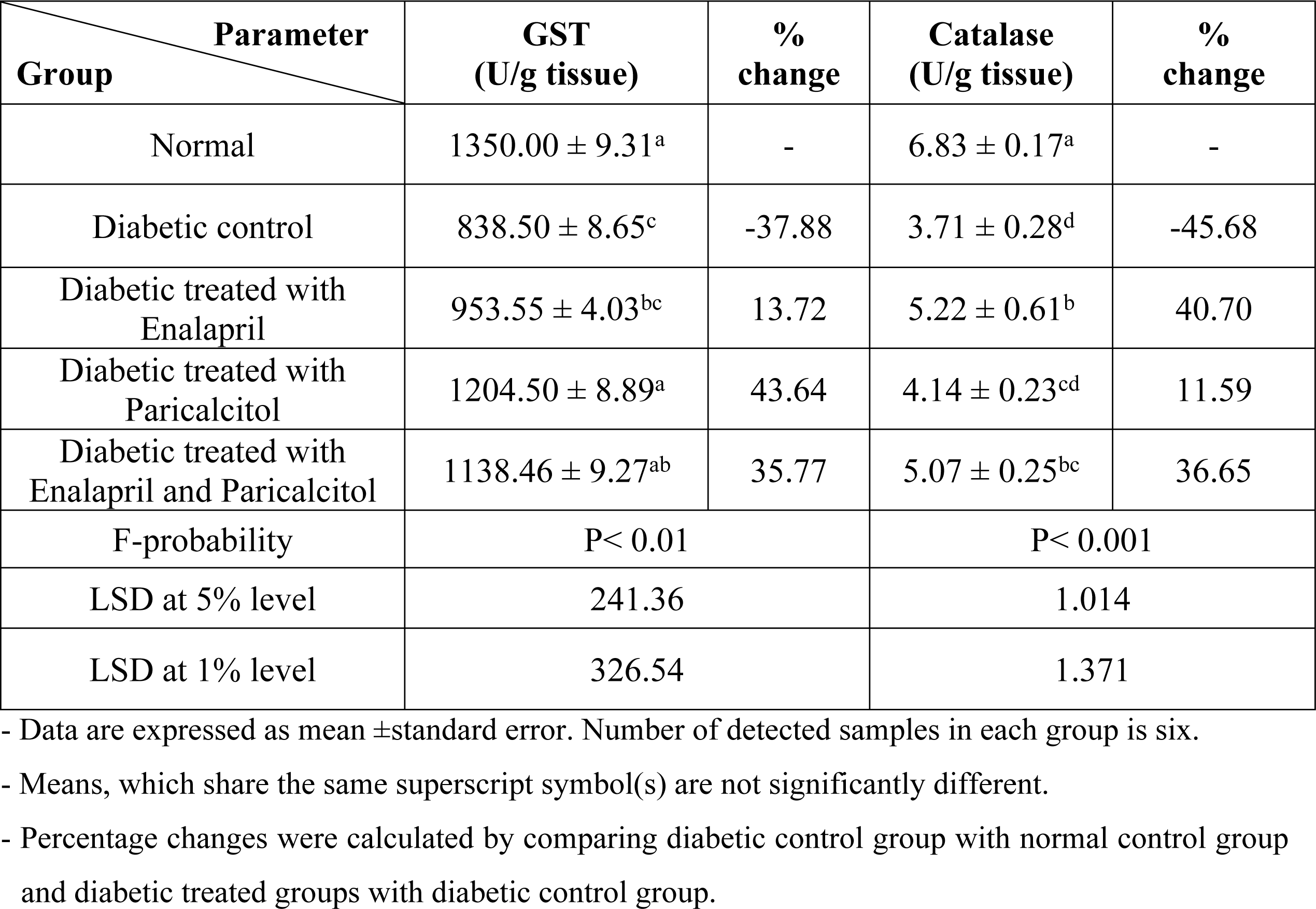
Effects of enalapril and paricalcitol on kidney GST and catalase activities in diabetic rats.

On the other hand, the elevated fasting and postprandial blood glucose levels in diabetic rats were significantly improved (p<0.01; LSD) as a result of treatments with enalapril, paricalcitol and their combination. The diabetic group treated with enalapril and paricalcitol together exhibited the most potent effects in ameliorating the elevated serum fasting and postprandial glucose level; the recorded percentage changes were −63.55 and −60.37% respectively (Table 5). The serum insulin and C-peptide levels were significantly decreased in diabetic rats recording percentage changes of −76.98 and −83.64% respectively. The treatment with enalapril and/or paricalcitol successfully led to a significant increase of the lowered serum insulin and C-peptide levels. The treatment with enalapril was the most potent in increasing serum insulin and C-peptide levels recording percentage increases of 517.05 and 83.64% respectively (Table 6).

**Table 5:** Effects of enalapril and paricalcitol on serum fasting and postprandial glucose levels in diabetic rats.

| Parameter<br>Group | Fasting blood<br>glucose<br>(mg/dl) | %<br>change | Postprandial<br>blood glucose<br>(mg/dl) | %<br>change |
| --- | --- | --- | --- | --- |
| Normal | 70.01 ± 5.91<sup>d</sup> | - | 98.00 ± 5.34<sup>d</sup> | - |
| Diabetic control | 229.40 ± 12.56<sup>a</sup> | 231.26 | 255.75 ± 7.67<sup>a</sup> | 160.96 |
| Diabetic treated with<br>Enalapril | 135.25 ± 6.91<sup>b</sup> | -41.04 | 190.66 ± 16.10<sup>b</sup> | -25.45 |
| Diabetic treated with<br>Paricalcitol | 103.60 ± 5.09<sup>c</sup> | -54.83 | 157.41 ± 11.93<sup>c</sup> | -38.45 |
| Diabetic treated with<br>Enalapril and Paricalcitol | 83.6 ± 3.66<sup>cd</sup> | -63.55 | 101.33 ± 2.56<sup>d</sup> | -60.37 |
| F-probability | P< 0.001 |  | P< 0.001 |  |
| LSD at 5% level | 21.806 |  | 29.009 |  |
| LSD at 1% level | 29.502 |  | 39.247 |  |

- Data are expressed as mean ± standard error. Number of detected samples in each group is six. - Means, which share the same superscript symbol(s) are not significantly different. - Percentage changes were calculated by comparing diabetic control group with normal control group and diabetic treated groups with diabetic control group.

**Table 6:** Effects of enalapril and paricalcitol on serum insulin and C-peptide levels in diabetic rats.

| Parameter<br>Group | Insulin<br>(ng/ml) | %<br>change | C-peptide<br>(ng/ml) | %<br>change |
| --- | --- | --- | --- | --- |
| Normal | 2.633 ± 0.111<sup>a</sup> | - | 4.733 ± 0.264<sup>a</sup> | - |
| Diabetic control | 0.606 ± 0.189<sup>c</sup> | -76.98 | 0.774 ± 0.109<sup>c</sup> | -83.64 |
| Diabetic treated with<br>Enalapril | 1.833 ± 0.152<sup>b</sup> | 202.47 | 4.776 ± 0.097<sup>a</sup> | 517.05 |
| Diabetic treated with<br>Paricalcitol | 1.413 ± 0.195<sup>b</sup> | 133.16 | 4.200 ± 0.146<sup>a</sup> | 442.63 |
| Diabetic treated with<br>Enalapril and Paricalcitol | 1.640 ± 0.204<sup>b</sup> | 170.62 | 3.233 ± 0.427<sup>b</sup> | 317.70 |
| F-probability | P< 0.001 |  | P< 0.001 |  |
| LSD at 5% level | 0.5063 |  | 0.7084 |  |
| LSD at 1% level | 0.6850 |  | 0.9854 |  |

- Data are expressed as mean ± standard error. Number of detected samples in each group is six. - Means, which share the same superscript symbol(s) are not significantly different.

Histopathological examination of kidney tissues of diabetic group showed severe necrosis of epithelial cells lining renal tubules, intense congestion of glomerular tuft (Fig 1; Photomicrograph B) and intertubular hemorrhage as compared with kidney histological structure of normal group (Fig 1; Photomicrograph A). The kidney section of diabetic group treated with enalapril showed no histopathological changes (Fig 1; Photomicrograph C) while mild congestion of glomerular tuft and intense congestion of blood vessels were observed in diabetic rats treated with paricalcitol. Moreover, although there are inflammatory cells in diabetic group treated with a combination of both treatments, most of tubules appeared with normal intact architecture (Fig 1; Photomicrograph D).

**Fig 1:**
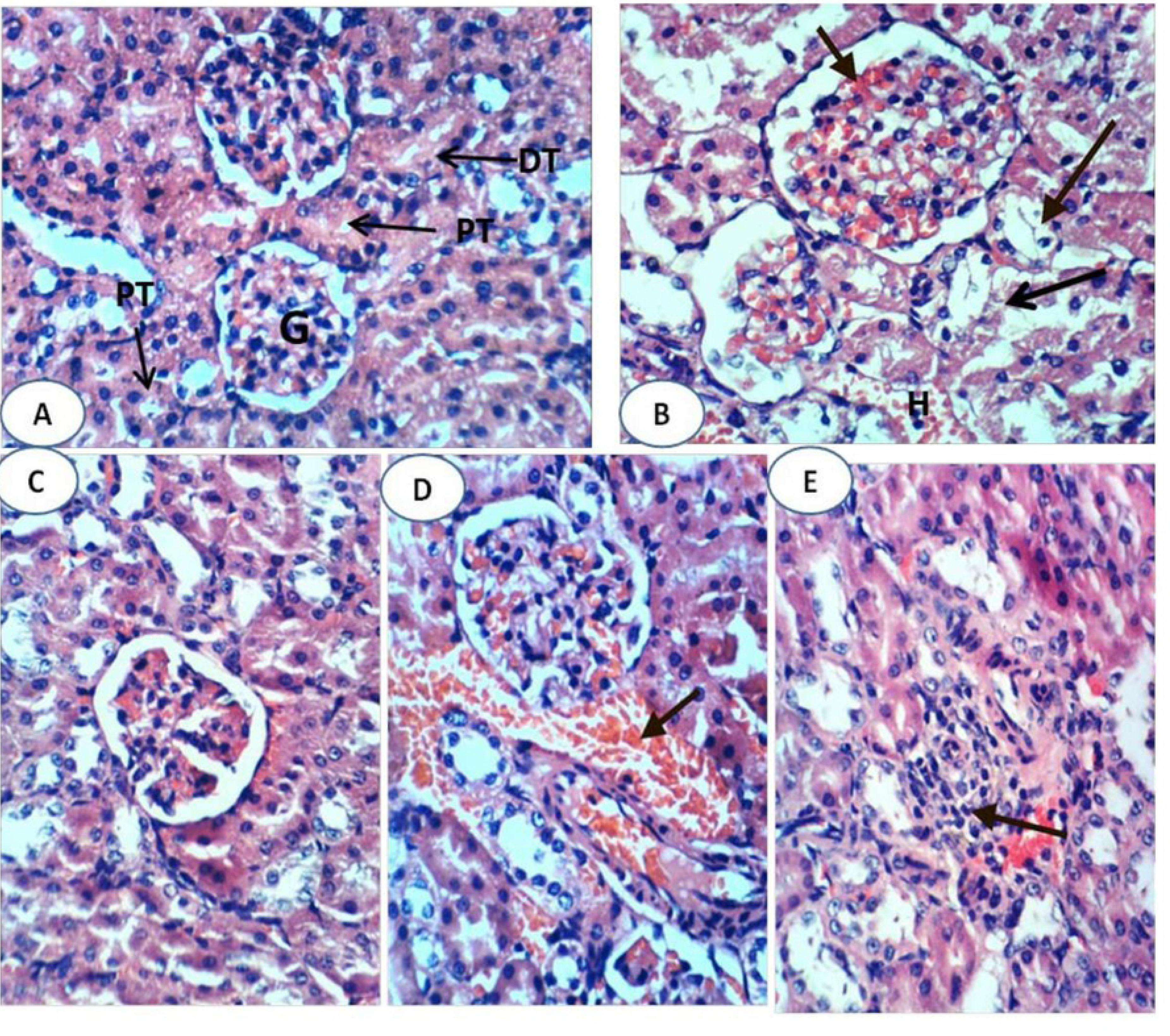
Photomicrographs of hematoxylin and eosin (H & E) stained sections. Photomicrograph A is showing normal kidney histological structure with normal glomerulus (G), normal proximal tubule (PT) and normal distal tubule (DT). Photomicrograph B is showing severe diffuse tubular necrosis (long arrow), severe congestion of glomerular tuft (short arrow) and intertubular hemorrhage (H) in diabetic group. Photomicrograph C is showing normal histological structure in diabetic group treated with enalapril. Photomicrograph D is showing congestion of blood vessel (arrow) in diabetic group treated with paricalcitol. Photomicrograph E is showing focal necrosis and inflammatory cell infiltration (arrow) in diabetic rats treated with both enalapril and paricalcitol (E). (H&E; 400X)

Immunohistochemical studies revealed no or week expression of kidney TNF-α in normal control group (Figure 2; Photomicrograph A) and in diabetic groups treated with enalapril (Figure 2; Photomicrograph C), paricalcitol (Figure 2; Photomicrograph D) and their combination (Figure 2; Photomicrograph E) while intense expression in diabetic control group (Figure 2; Photomicrograph B). Kidney P53 exhibited negative expression in normal group (Figure 3; Photomicrograph A), diabetic groups treated with enalapril (Figure 3; Photomicrograph C) and its combination with paricalcitol (Figure 3; Photomicrograph E) while it showed moderate expression in diabetic rats treated with paricalcitol (Figure 3; Photomicrograph D) and strong positive expression in diabetic control group (Figure 3; Photomicrograph B). Kidney caspase-3 exhibited negative expression in normal group (Figure 4; Photomicrograph A), diabetic group treated with enalapril (Figure 4; Photomicrograph C) and its combination with paricalcitol (Figure 4; Photomicrograph E) while it showed strong positive expression in diabetic control (Figure 4; Photomicrograph B) and mild expression in diabetic group treated with paricalcitol (Figure 4; Photomicrograph D). Conversely, Kidney Bcl-2 showed weak expression in diabetic control group (Figure 5; Photomicrograph B) and strong positive expression in normal group (Figure 5; Photomicrograph A), diabetic groups treated with enalapril (Figure 5; Photomicrograph C), paricalcitol (Figure 5; Photomicrograph D) and their combination (Figure 5; Photomicrograph E).

**Fig 2:**
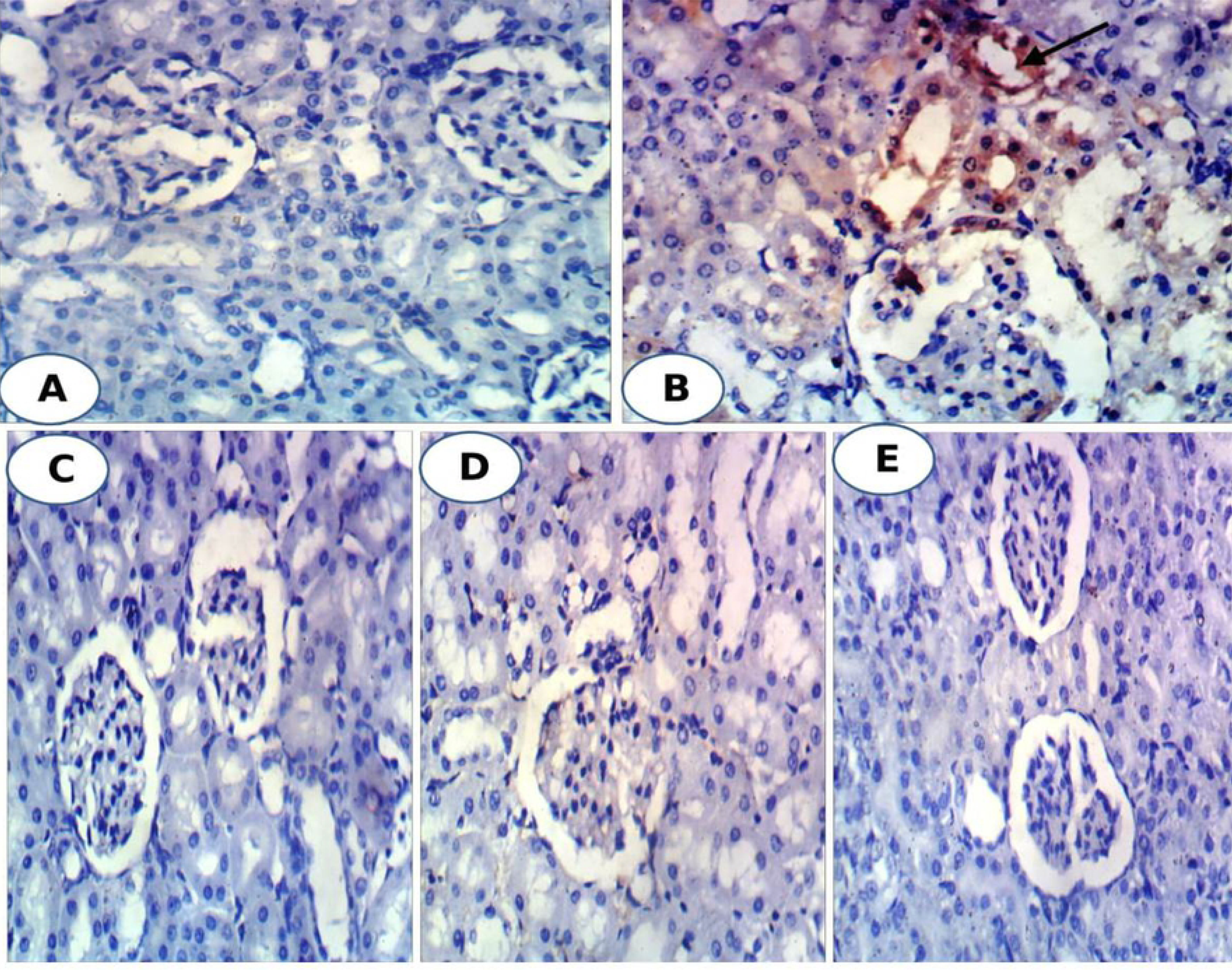
Photomicrographs of immunohistochemical staining of TNF-α in kidney tissues showing negative expression in normal group (A), diabetic groups treated with enalapril (C), paricalcitol (D) and their combination (E) while strong positive expression in diabetic control (B). (400X)

**Fig 3:**
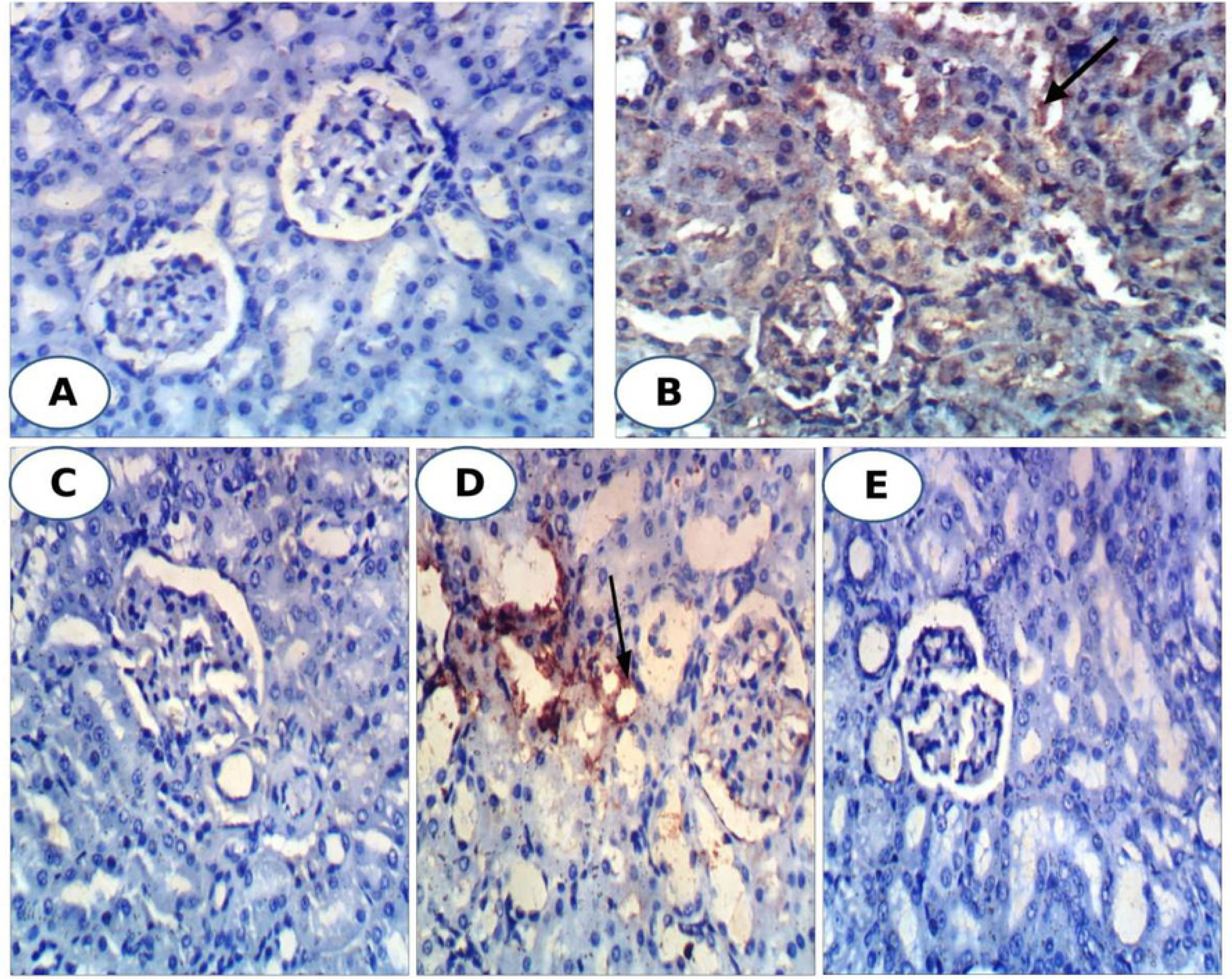
Photomicrographs of immunohistochemical staining of P53 in kidney tissues showing negative expression in normal group (A), diabetic groups treated with enalapril (C) and its combination with paricalcitol (E) while moderate expression in diabetic rats treated with paricalcitol (D) and strong positive expression in diabetic control group (B). (400X)

**Fig 4:**
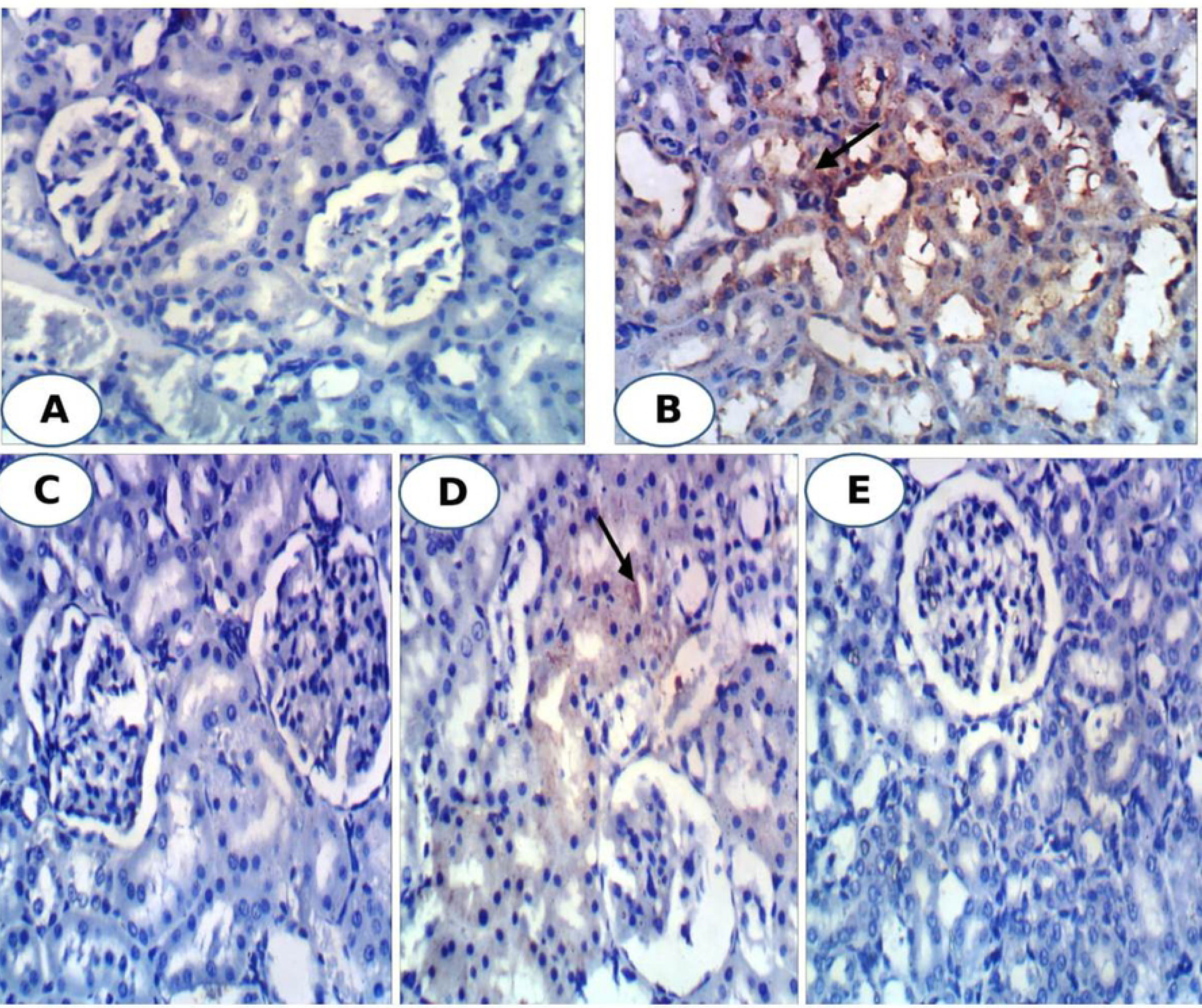
Photomicrographs of immunohistochemical staining of caspase 3 in kidney tissues showing negative expression in normal group (A), diabetic group treated with enalapril (C) and its combination with paricalcitol (E) while strong positive expression in diabetic control (B) and mild expression in diabetic group treated with paricalcitol (D). (400X)

**Fig 5:**
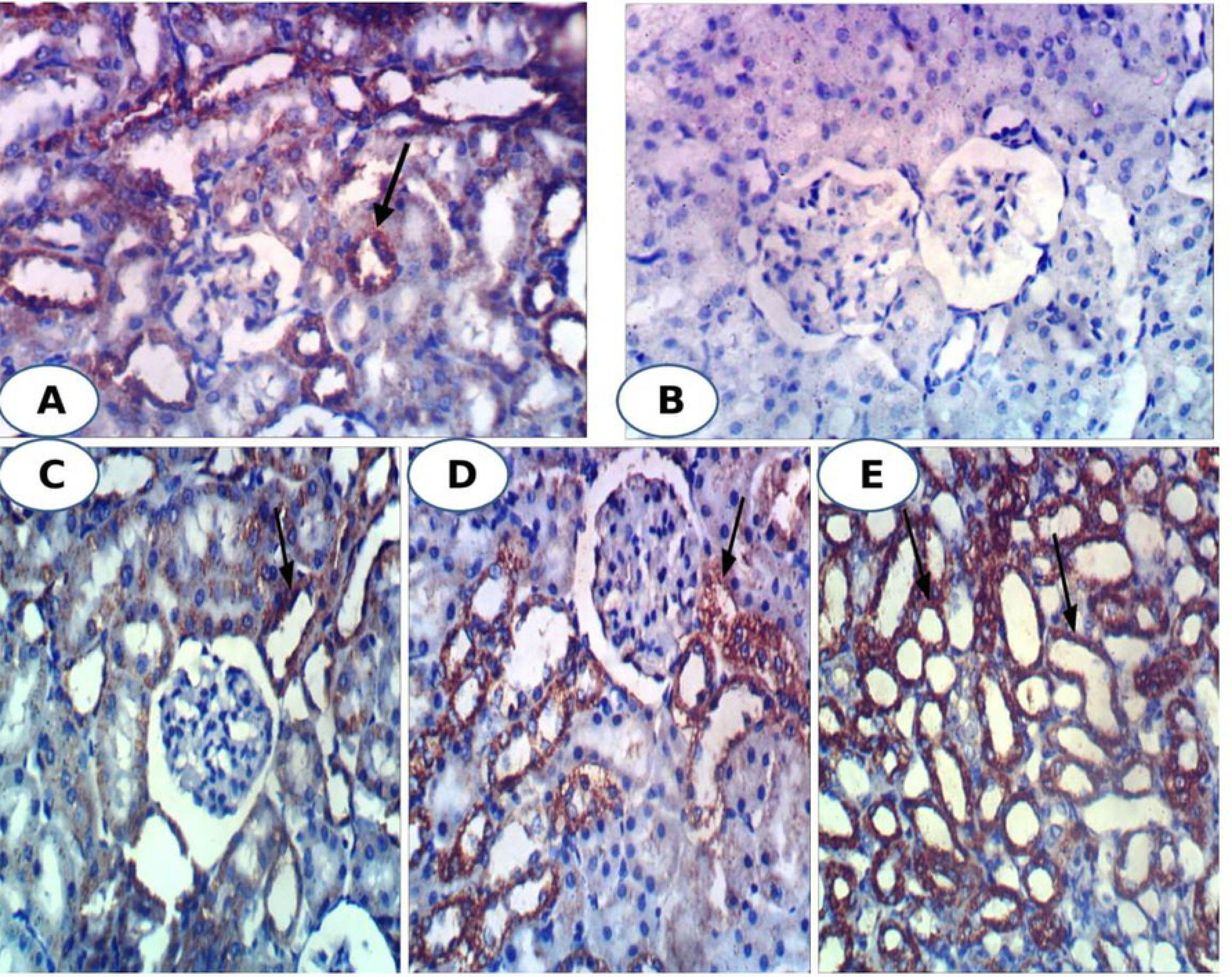
Photomicrographs of immunohistochemical staining of Bcl-2 in kidney tissues showing weak expression in diabetic control group (B) and strong positive expression in normal group (A), diabetic groups treated with enalapril (C), paricalcitol (D) and their combination (E). (400X)

The pancreas sections of normal rats showed normal islet histological architecture and integrity in (Figure 6; Photomicrograph A). The islets of Langerhans of diabetic rats exhibited necrotic and destructive changes of islets cells and decrease in the size of islets (Figure 6; Photomicrograph B) as compared with normal islets. The diabetic rats treated with enalapril showed marked increase of islet size but still vacuolations are present (Figure 6; Photomicrograph C). The diabetic rats treated with paricalcitol showed intact islet with normal architecture (Figure 6; Photomicrograph D). The diabetic rats treated with a combination of enalapril and paricalcitol exhibited relatively large-sized islets with highly active divided cells (Figure 6; Photomicrograph E).

**Fig 6:**
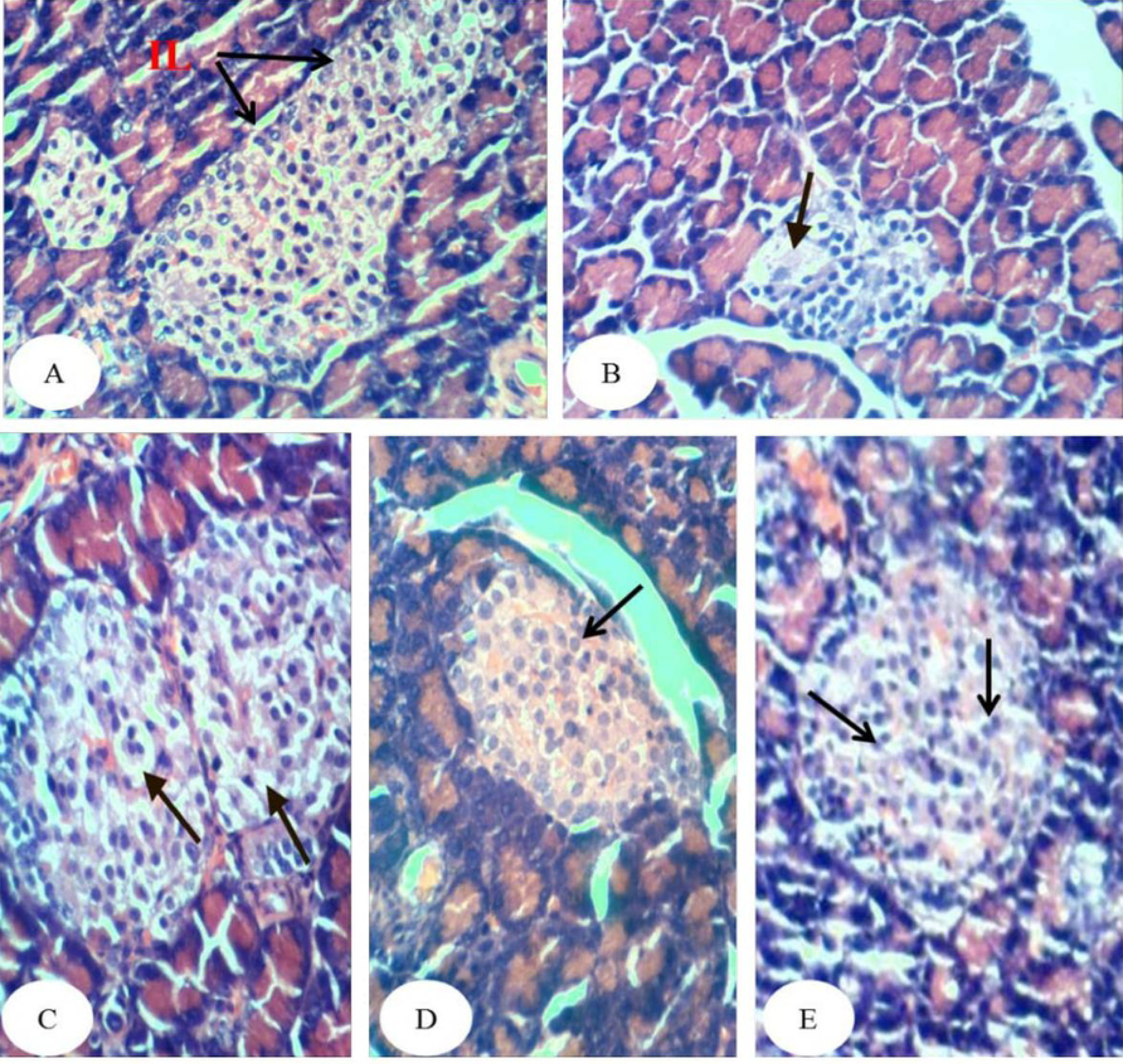
Photomicrographs of pancreas sections showing no histopathological changes in normal group (A), necrosis (arrow) and decrease in the size of islets of diabetic control group (B), marked increase of islet size and vacuolations (arrow) in diabetic group treated with enalapril (C), intact islet in diabetic group treated with paricalcitol (D) and relatively large-sized islets with highly active divided cells in diabetic group treated with a combination of enalapril and paricalcitol (E). (H&E; 400X)

## Discussion

Diabetic nephropathy, a severe complication of diabetes mellitus, is the most common cause of end stage renal failure. About 15–25% of type 1 diabetes patients and 30–40% of patients with type 2 diabetes suffer from diabetic nephropathy [4]. In this context, the model of type 1 diabetes, rat STZ-induced diabetes model, is used in the present study to investigate the pathogenesis of diabetic nephropathy [24]. Streptozotocin is an agent of choice to induce experimental diabetes mellitus due to its ability to induce specific necrosis of the pancreatic beta cells that results in degranulation and loss of capacity to secrete insulin [25], thereby leading to hyperglycemia and diabetic complications such as nephropathy [24]. As indicated in the present study, STZ induces a significant increase fasting and post-prandial serum glucose levels, a significant depletion in serum insulin and C-peptide levels, decrease in the size of islets and necrotic changes in the islet cells and subsequent kidney dysfunction.

In the present study, the STZ-induced diabetic rats exhibited impairment in kidney function that was manifested by a significant elevation of serum urea, uric acid, creatinine sodium and potassium levels as well as derangement in kidney histological architecture and integrity which was marked by severe glomerular congestion, tubular necrosis and intertubular hemorrhage. These results are in accordance with Ahmed [26] who reported a significant increase in serum urea, uric acid and creatinine levels tandem to severe hyperemia in the glomerular tufts, intertubular hyperemia and degenerative changes in the epithelium cells lining the renal tubules in STZ-induced diabetic rats. The damaging effects of STZ on kidney may be attributed to the increased oxidative stress and the attenuated antioxidant defense system. The present study supports this attribution since the STZ-induced diabetic rats exhibited a significant elevation in the kidney LPO and a significant decrease in kidney GSH content and suppression of GST and catalase activities. In agreement with these results, de Brito Amaral *et al*. [27] revealed a significant increase in kidney LPO in sedentary and trained STZ-induced diabetic rats as compared with normal control. Moreover, Ziamajidi *et al*. [28] confirmed the elevation of LPO and total oxidative stress in nicotinamide (NA)/STZ-induced diabetic rats. In our opinion, it can be stated that the persistent hyperglycemia in diabetes mellitus leads to increased production of reactive oxygen species (ROS) which are involved in the etiology of several diabetic complications including diabetic nephropathy. The reactive oxygen species deplete the antioxidant defenses of the cell thus making it more susceptible to oxidative damage [29]. ROS further target lipid, DNA and protein leading to their oxidation which further lead to changes in cellular structure and function [30].

In addition to the role of oxidative stress in inducing the renopathy, the inflammation and apoptosis may have an important role in eliciting kidney dysfunction and histological deteriorations since the STZ-induced diabetic rats, in the present study, exhibited a remarkable increase in the expression of immunohistochemically-detected pro-inflammatory cytokine, TNF-α and apoptotic markers including P53 and caspase-3 as well as a decrease in the anti-apoptotic markers Bcl-2. These results are in concurrence with the previous study of Pradeep and Srinivasan [31] who demonstrated an increased apoptotic mediator, Bax and decrease in Bcl-2, detected by immunoflourscence and immunohischemical techniques, in kidney of STZ-induced diabetic rats.

The treatment of diabetic rats with enalapril and/or paricalcitol, in the present study, resulted in a marked improvement of kidney function represented by a significant decrease in the elevated serum urea, uric acid and creatinine levels along with a remarkable amelioration of the deteriorated kidney histological changes. These ameliorations in kidney function and histological architecture and integrity are associated with the improvements in the glycemic state, serum insulin and C-peptide levels, islets histological changes, kidney oxidative stress and antioxidant defense system, kidney TNF-α as pro-inflammatory cytokine, and kidney apoptotic (P53 and caspase-3) and anti-apoptotic (Bcl-2) markers.

The pathogenesis of renal injury included complex pathway crosstalk contributing to increased oxidative stress and inflammation, as well as renal tubular apoptosis, during the disease course [32]. As STZ diabetes-induced oxidative stress, local inflammation and tubular apoptosis were implicated in the pathogenesis of renal dysfunction as confirmed in this study and previous publications [27,28,31], the improvement effects of enalapril and/or paricalcitol on these processes in STZ-induced diabetic rats may be all involved to alleviate the kidney function and kidney structural integrity. In this way, enalapril, a non-sulfhydryl ACEI, has shown renoprotective effect in various animal models including diabetic nephropathy animal model [33]. Clinically, enalapril exhibited renoprotective effect in patients with chronic kidney disease, diabetic nephropathy, and hypertension after renal transplantation [34,35]. On the other hand, paricalcitol, which is an active non-hypercalcemic vitamin D analogue, displays analogous biological activity, with scarcer adverse effects and amplified tolerance, as matched to those of vitamin D. In this regard, apoptosis inhibition was reported in case of STZ diabetic rats after observing higher levels of insulin and C-peptide in diabetic group receiving active vitamin D3 [36]. Vitamin D may also mitigate kidney damage by overwhelming fibrosis, inflammation, and apoptosis, by hindering multiple pathways including RAAS, the nuclear factor-κB (NF-κB), the transforming growth factor-β (TGF-β)/Smad, and the Wnt/β-catenin signaling pathways [37,38,39].

The increased LPO in the renal tissue observed in diabetic rats, in present study, was significantly suppressed as a result of treatment with enalapril and/or paricalcitol while the lowered GSH content as well as catalase and GST activities were increased, thus suggesting the antioxidant capacity of paricalcitol and enalapril. The decreased LPO after treatment with enalapril and/or paricalcitol, in the present study, could be a result of the increase in the above mentioned antioxidant components which scavenge excess ROS before extreme oxidation can occur. In agreement with the present study, it was reported that enalapril treatment, on account of its antioxidant properties, reduced oxidative stress in the kidney tissues of spontaneously hypertensive rats [40] and streptozotocin-induced diabetic rats [33]. The present results are also in consistence with a previous study of the effect of paricalcitol in hemodialysis patients that had revealed a similar effect of paricalcitol on catalase enzyme [15].

Consequentially, glomerular and tubular TNF-α, in the present study, showed increased expression in diabetic kidney and exhibited a decreased expression in diabetic rats treated with enalapril and/or paricalcitol reflecting the anti-inflammatory effect of these drugs. These results are in concordance with Navarro *et al*. [41] who stated that enalapril administration nearly completely abolished the increase in renal TNF-α messenger RNA expression and reduced urinary albumin excretion (UAE) in STZ-induced diabetic rats reflecting the role of anti-inflammatory effect in in the improvement of kidney function. Those authors also found that blockade of the renin-angiotensin system (RAS) by enalapril prevent the enhanced expression of TNF-α suggesting the possible regulatory role of RAS on renal inflammatory status. The present results also go parallel with Izquierdo *et al*. [15] who found that after paricalcitol treatment of patients with renal disease, levels of the inflammatory markers CRP, TNF-α, IL-6 and IL-18 were significantly reduced in serum and the level of anti-inflammatory cytokine IL-10 was increased. The anti-inflammatory possessions of active vitamin D and its analogues (such as paricalcitol) as well as enalapril may be endorsed to their ability to overwhelm the NF-κB pathway, a key transcription factor that is supposed to facilitate acute and chronic inflammation by regulating gene expression of cytokines and chemokines (including interleukin-6 and tumor necrosis factor-α) [42]. NF-κB activation associated with increased ROS generation is pivotal in the consequent expression of proinflammatory cytokines like TNF-α. These chemokines may then facilitate migration and infiltration of inflammatory cell and a secondary wave of ROS generation, and further amplify the inflammatory cascade and injury [43]. In another way, paricalcitol weakens renin and angiotensin II expression in VDR knockout mice [44], telling that paricalcitol inhibits renal inflammation by overwhelming the RAAS, as angiotensin II is a known proinflammatory stimulus. Taken together, these findings indicate that enalapril and vitamin D analogue, paricalcitol, might be useful for treating inflammatory kidney diseases.

In trial to elucidate the role of apoptosis in the pathogenesis of diabetic nephropathy and in the ameliorative effects of enalapril and paricalcitol, renal apoptotic proteins including P53 and caspase-3, and anti-apoptotic protein, Bcl-2, were immunohistochemically detected. The present study indicated that the expression of tubular P53 and caspase-3 were remarkably increased in diabetic rats and were decreased as a result of treatment with enalapril and/or paricalcitol. The effect of enalapril alone and enalapril concomitant with paricalcitol seemed to be more potent in decreasing the expression of tubular P53 and caspase-3 than the effect of paricalcitol alone. The tubular expression of anti-apoptotic protein Bcl-2 exhibited a reverse pattern of changes with P53 and caspase-3. All these observations indicate a protective effect of paricalcitol and enalapril on STZ-induced renal tubular apoptosis probably *via* modifying expression of P53, Bcl-2 family proteins and subsequent caspase-3.

The p53 protein corresponds to a number of processes associated with the life and death of the cell. It regulates the repair of cellular DNA and encourages apoptosis when the damage of the gene is too serious and it is difficult to repair [45]. It has also been established that the p53 protein, as a consequence of stress factors, crosses into the mitochondria and triggers the expression of pro-apoptotic genes, for example Puma, Bax, Apaf-1, Noxa. In addition, it inhibits the expression of anti-apoptotic genes, like those of the Bcl-2 family (Bcl-2, Bcl-X, Bcl-in, Mcl-1); this evidenced was confirmed in the present investigation since the tubular expression of anti-apoptotic protein Bcl-2 exhibited a reverse pattern of changes with P53. The p53 protein together with these pro-apoptotic proteins, are moved into the mitochondria where they encourage an upsurge in the permeability of mitochondrial membranes and the release of cytochrome C, which attaches to the apoptotic protease activating factor 1 (Apaf-1) and with the caspase-9 proenzyme, to generate the complex called the “apoptosome”. The apoptosome, consecutively, produces activation of caspase-9. The later consequently stimulates the caspase-3 proenzyme activation to the protease stage, which then sticks to the effector caspase group. These caspases induce intracellular protein lysis and morphological distinctive changes of apoptosis [45]. It was reported that apoptosis could aggravate the pathogenesis of nephrotoxicity *via* caspase-3 expression [46]. Mitochondrial oncogene product, Bcl-2, prevented caspase-3 activation during a variety of proapoptotic conditions [47].

The antiapoptotic effect of enalapril observed in the current study was also in accordance with the study of Rani *et al*. [48] who revealed that pretreatment with enalapril dose dependently restored the cisplatin-induced apoptosis towards normal as it reduced Bax, caspase-3, cytochrome C, and p53 expressions with concomitant increase in the Bcl-2 expression thus resulting in anti-apoptotic potential. The antiapoptotic effects of paricalcitol was also reported. A previous study showed that paricalcitol reduces the increased expression of phospho-p53 which recruits apoptotic processes in a cisplatin-induced rat model [49]. Moreover, it was reported that an increased Bax/Bcl-2 ratio and the cleaved form of caspase-3, which are apoptotic markers, are reversed by paricalcitol treatment in gentamicin-induced kidney injury [50]. Thus, the enalapril and paricalcitol have anti-apoptotic effects and p53, caspase-3 and Bcl-2 may have crucial role to mediate these effects.

## Conclusions

The study concluded that paricalcitol and/or enalapril potentially protect against STZ-induced diabetic renopathy via their potencies to improve the diabetic condition, suppress the oxidative stress, enhance the antioxidant defense system and decrease the apoptosis through attenuating the renal expression of P53 and caspase-3 and enhancing the expression of Bcl-2.

## Acknowledgement

The authors thank Prof. Dr. Kawkab Abdel Aziz Ahmed, Professor of Histopathology, Pathology Department, Faculty of Veterinary Medicine, Cairo University, Egypt and Prof. Dr. Rasha R. Ahmed, Professor of Molecular Cell Biology, Cell Biology, Histology and Genetics Division, Zoology Department, Faculty of Science, Beni-Suef University, Beni-Suef, Egypt for help in examining and reading the histological and immunohistochemical stained sections

## Data availability

The datasets generated and/or analyzed during the current study are available from the corresponding author on reasonable request.

## Author contributions

OMA, TMA and MAA designed the study and wrote and edited the manuscript. OMA and MAA performed the experiments and analyzed the data. OMA, TMA, MAA and AAE revised and approved the manuscript.

